# Modelling the transport of fluid through heterogeneous, whole tumours *in silico*

**DOI:** 10.1101/512236

**Authors:** Paul W. Sweeney, Angela d’Esposito, Simon Walker-Samuel, Rebecca J. Shipley

**Affiliations:** Mechanical Engineering, University College London, London, United Kingdom; Centre for Advanced Biomedical Engineering, University College London, London, United Kingdom

## Abstract

It is critically important to understand and predict fluid transport within both physiological and pathological tissues in order to develop effective treatment strategies. Recent advances in high-resolution optical imaging allow the acquisition of whole tumour vascular networks which can be used to parameterise computational models to predict the fluid dynamics at all length scales across the tissue. This enables hypothesis testing around the role of the tumour microenvironment in determining transport characteristics, which would otherwise be unavailable using traditional experiments.

In this study, we present a novel computational framework for the efficient simulation of vascular blood flow and interstitial fluid transport based on complete three-dimensional, whole tumour vasculature obtained using high-resolution optical imaging. This framework comprises a Poiseuille flow model which simulates vascular blood flow within the vessel network, coupled via point sources of flux to a porous medium model describing interstitial fluid transport. We develop a computational algorithm for prescription of network boundary conditions and validation of tissue-scale fluid transport against measured *in vivo* perfusion data acquired using biomedical imaging tools. We present simulations of the model on orthoptic murine glioma and human colorectal carcinoma xenograft data (GL261 and LS147T, respectively), and perform sensitivity analysis on key unknown parameters relating to the tissue microenvironment, to understand their impact in predicting vascular and interstitial flow. Finally, we simulate radially varying vascular normalisation in a LS147T tumour and hypothesise that uniform normalisation is required to lower tumour interstitial fluid pressure.

Our computational framework permits predictions of whole tumour fluid dynamics which incorporate the inherent architectural heterogeneities appearing at the micron-scale, and outputs three-dimensional spatial maps detailing these flow properties from micro to macro length scales. This provides vital information on the tumour microenvironment which could enable the design and delivery of future anti-cancer therapies.

**Author summary:** The structure of tumours varies widely, with dense and chaotically-formed networks of blood vessels that differ between each individual tumour and even between different regions of the same tumour. This atypical environment can inhibit the delivery of anti-cancer therapies. Computational tools are urgently required which incorporate micron-scale tumour biomechanics to predict tissue-scale fluid dynamics, and consequently the efficacy of cancer therapies.

We have developed a computational framework which integrates the complex tumour vascular architecture to predict fluid transport across all lengths scales in whole tumours. This enables computationally efficient hypothesis testing of cancer therapies which manipulate the tumour microenvironment in order to improve drug delivery to tumours.

## Introduction

Architectural heterogeneities in cancerous tissue limit the delivery of anti-cancer drugs by inhibiting their ability to circumnavigate the entire tumour to all cancerous cells [1]. In solid tumours, drug penetration to the tumour core is hindered by physiological barriers which can limit the delivery of targeted agents, with penetration exacerbated by the size of the agent [1–5]. Consequently, preclinical tools which provide a better understanding of therapy interactions within the tumour microenvironment are urgently required in order to increase treatment efficacy. In *silico* modelling is one such tool which can meet this need by testing novel therapeutic strategies at a much faster rate and much cheaper cost than preclinical experimentation [6].

For a systemically-administered agent to effectively target diseased tissue, it must travel from the site of injection to the site of disease, whilst minimally interacting with normal tissues and not degrading [7]. This is difficult to achieve in tumours since atypical endothelial proliferation of tumour vasculature leads to spatial variations in vascular density and branching patterns, distorted and enlarged vessels, and a highly convoluted network topology [8–10]. Further, vascular permeability is heightened and heterogeneous and so these immature blood vessels are generally leakier than those in normal tissue [3, 11].

The irregular microenivronment is typically characterised by hypoxia, acidosis and elevated interstitial fluid pressure (IFP) [12–14], which drive both tumour vascular proliferation and resistance to therapy [15]. Here, drug delivery may be hindered by the atypical nature of the tumour interstitium. The extracellular matrix (ECM) consists of a cross-linked dense network of collagen and elastin fibres, far denser than usually seen in normal tissue [16]. A denser matrix can result in reduced delivery of oxygen and nutrients, as well as providing significant resistance to the advection and diffusion of therapeutic particles [1], since key determinants of intratumoural fluid and mass delivery include tissue hydraulic conductivity and vascular compliance [17]. Several therapeutic interventions have sought to limit the effects of these physical barriers by manipulating the microenvironment to enhance the delivery of macromolecular agents [16, 18]. For example, normalising the tumour vasculature to reduce vessel permeability thereby increasing drug penetration [12]; and manipulating the connective tissue, and therefore interstitial hydraulic conductivity, using a platelet-derived growth factor (PDGF) antagonist to reduce tumour IFP [19].

Heterogeneities in the underlying morphology of tumours, such as vessel diameters and lengths, and inter-branch distance, exist across individual tumours and tumour cell-lines [20]. These variations in tumour architecture lead to spatial variability in drug efficacy, which complicate efforts to design effective treatment strategies [7]. Experimental efforts have been made to understand the effects of tumour heterogeneity on fluid interactions across tumours, for example, wick-in-needle has been used to measure IFP across tumours [21–23]. However, this method disturbs the local microenvironment and only provides IFP measurement at individual locations. Non-invasive methods have also been developed to estimate tumour IFP [24, 25]. For example, convection-MRI, which, with further validation, presents an opportunity to measure low-velocity flow in tumours, and to assess therapeutic response [26]. However, these methods fail to capture full spatial maps of flow at the micron-scale which are crucial to understanding how the combined intra- and extravascular spatial flow heterogeneities occurring at the scale of blood vessels affects the macro-scale flow dynamics and consequent delivery of drugs within a solid tumour. Biomedical imaging complemented by in *silico* methods provides scope to provide such detail.

Mathematical models have been used to investigate the tumour microenvironment and have provided detailed insights which may otherwise be unavailable experimentally. Seminal models have indicated that a leaky tumour vasculature induces elevated IFP, reduced fluid penetration into the interstitium [14, 27], and a non-uniform distribution of drug delivered to solid tumours [2, 3, 11]. Further, they have defined conventional IFP profiles in tumours - a uniform pressure at the core, with a large decreasing gradient towards the periphery. However, these models generally average over the tumour vasculature and so fail to capture the micron-scale flow dynamics; and they assign a fixed pressure boundary condition on the periphery of the tumour which may artificial induce these conventional IFP profiles. Subsequent studies have incorporated the spatially heterogeneous effects of tumour vasculature using computer-generated synthetic networks which retain key features of tumour vascular architecture [28–34], or by integrating spatial variations in vascular permeability parametrised against *in vivo* experimentation [35, 36]. However, to date, *in silico* models have lacked realistic, high-resolution data on whole tumour vascular architecture to both parametrise and validate computational models [6].

Recent advances in *ex vivo* optical imaging of cleared tissue specimens have enabled large samples (up to 2 cm^3^ with > 10^5^ blood vessels) to be imaged in three-dimensions, at resolutions down to a few microns [37]. We have developed a platform called REANIMATE (<u>REA</u>listic <u>N</u>umerical <u>I</u>mage-based <u>M</u>odelling of biologic<u>AL</u> <u>T</u>issue substrat<u>E</u>s) which combines optical imaging of cleared tissue with mathematical modelling and *in vivo* imaging, within a unified framework, to generate quantitative, testable predictions regarding tumour transport [38]. The platform uses high-resolution imaging data from large, intact, optically-cleared tissue samples to make *in silico* predictions of blood flow, vascular exchange and interstitial transport. REANIMATE enables new hypotheses to be generated and tested in a manner that would be challenging or impossible in a conventional experimental setting. As a proof-of-principle, we have previously used REANIMATE to explore the impact of vascular network topology on fluid and therapy delivery, focusing on delivery of a vascular disrupting agent (Oxi4503) to two colorectal cell-lines (LS147T and SW1222) [38].

A vital component of REANIMATE is the simulation of fluid transport across cancerous tissue. We developed a computational model to efficiently simulate both intra- and extravascular fluid transport across large, discrete microvascular networks. Our model simulates Poiseuille flow through the vasculature using the optimisation scheme of Fry et al. [39], parametrised and validated against *in vivo* ASL-MRI data [38]. Following a similar Green’s function method for oxygen transport [40], the vascular component is coupled, via a discrete set of point sources of flux, to a Darcy model which simulates the effective fluid transport in the porous interstitium. A linear system is formed whereby only the source strengths need to be resolved, making it more computationally efficient compared to finite difference or element methods which require a spatial, numerically-discretised mesh [40].

In this study, we present this model in detail along with a description of its application to whole tumour vascular networks. We apply our model to an orthotopic murine glioma and a human colorectal carcinoma xenograft from the GL261 and LS147T cell-lines, respectively, and reproduce physiological conditions observed in literature. We then perform sensitivity analysis to the model parameters associated with transvascular fluid delivery, such as vascular hydraulic conductance and interstitial hydraulic conductivity, to explore the impact on the tumour IFP and IFV profiles. Subsequently, we present preliminary predictions of vascular normalisation to an LS147T network. These results present an example of how our mathematical model can be used to simulate the heterogeneous pharmacokinetics of drug therapies designed to alter the properties of the tumour microenvironment, and provide three-dimensional spatial maps detailing changes in flow characteristics which indicate the efficacy of such treatments.

## Materials and methods

### Acquisition and Processing of Real-world Tumour Datasets

Orthotopic murine gliomas and human colorectal carcinoma xenograft from the GL261 and LS147T cell-lines (n = 6 for each), respectively, were grown subcutaneously in 8 – 10 week old, female mice. Following 10 to 14 days of growth, *in vivo* ASL-MRI was performed on a subset of GL261 and LS147T tumours, from which a mean tumour perfusion of 130 ± 50 and 19 ± 8 ml/min/100g was measured [38], respectively. Following perfuse-fixation, tumours were harvested, optically cleared and imaged using OPT (Bioptonics, MRC Technologies, Edinburgh). All experiments were performed in accordance with the UK Home Office Animals Scientific Procedures Act 1986 and UK National Cancer Research Institute (NCRI) guidelines [41]. Full details of the experimental protocol is provided in d’Esposito et al. [38].

Whole-tumour blood vessels networks were segmented from the OPT data for both tumour types. A combination of three-dimensional Gaussian and Frangi filters were applied to the data to enhance vessel-like structures allowing for the segmentation of the blood vessels from the background (see Fig 1 a). Skeletonisation of these thresholded data was performed in Amira (Thermo Fisher Scientific, Hillsboro, OR), which also converted the data into graph format (interconnected network of nodes and segments with associated radii - see Fig 1 b). To ensure that vessel structures were accurately represented, three-dimensional networks were visually inspected against two-dimensional imaging slices for an accurate representation of vessel location and thickness. Full details of the validation can be found in the Supplementary Material of d’Esposito et al. [38].

**Fig 1.**
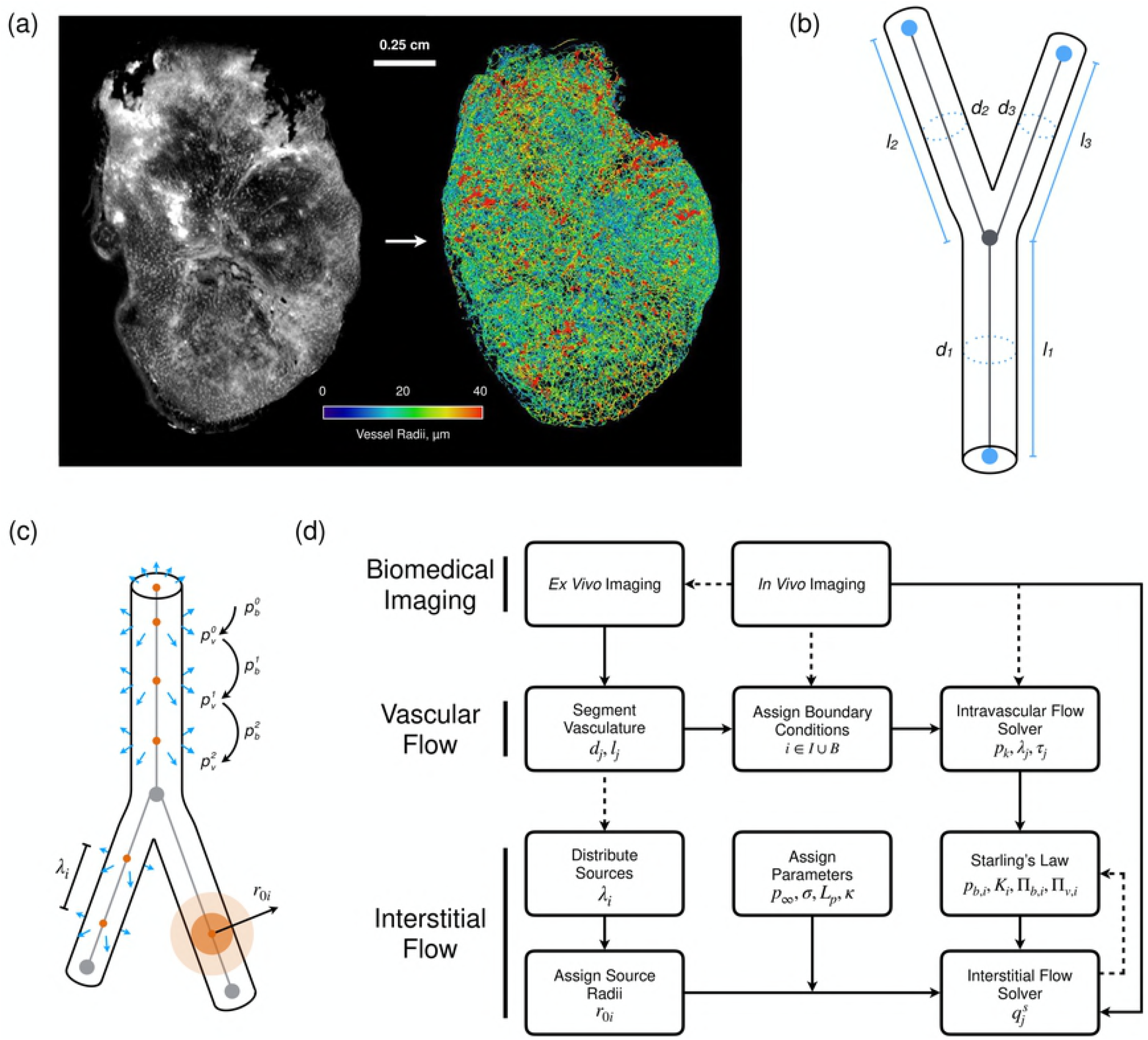
(a) An example of an SW1222 tumour vascular network enhanced using Frangi filters and extracted from the tumour image stack generated by OPT. (b) The skeletonised vasculature is then segmented into a series of interconnected nodes and vessel segments with known diameters, *d_i_* and lengths, *l_i_*, for *i* =1, 2,3. (c) A schematic of sensitivity analysis performed on the source parameters: 1) updating intravascular pressure *p_i,b_* for iterations i where *p*_0,*b*_ is the initial network pressure distribution approximated by the flow estimation algorithm; 2) the spacing λ_*i*_ between sources distribution across a branching vessel; and 3) the size of the source radius, *r*_0_. (d) A flow diagram of the computational framework. *In vivo* imaging is performed on vascularised tissue to obtain perfusion data (and literature values of vascular pressures when available) which are used to parameterise and validate the framework. *Ex vivo* imaging is performed on equivalent tissue samples to obtain data on the vascular architecture, including coordinates, vessel diameters and lengths, which are then used to parameterise the vascular flow model. Boundary conditions are assigned (see Fig 2 a and b) and network intravascular blood pressure is solved. Fluid sources of flux are distributed across the vasculature and assigned a radius equal to its corresponding vessel radius. Interstitial flow parameters are assigned and the model is coupled to the vascular flow compartment via Starling’s Law. Solved source strengths are used to update Starling’s law. This iterative scheme is terminated once predefined tolerances are reached.

In this study a GL261 and a LS147T tumour network were chosen from the d’Esposito et al. [38] datasets for *in silico* development and testing. Vessel diameters ranged from 17.9 ± 9.3 and 8.9 ± 2.8 *μ*m, with branching lengths of 68.7 ± 48.3 and 88.8 ± 49.4 *μ*m, respectively (see Table 1 and Fig 3). Vessel branching angles, inter-vessel distance, radii and tortuosity measures were consistent with data from previous studies that extracted vascular architectures using different methods [20, 38].

**Table 1.** Tumour vascular network statistics.

|  | GL261 | LS147T | Units |
| --- | --- | --- | --- |
| No. of Segments | 121,212 | 76,378 | - |
| No. of Nodes | 110,062 | 60,177 | - |
| No. of Boundary Nodes | 8,660 | 15,783 | - |
| Mean Vessel Diameter | $17.9 \pm 9.3$ | $8.9 \pm 2.8$ | $\mu\text{m}$ |
| Mean Branching Vessel Length | $68.7 \pm 48.3$ | $88.8 \pm 49.4$ | $\mu\text{m}$ |
| Tissue Dimensions | $4.3 \times 4.1 \times 4.6$ | $6.9 \times 5.2 \times 8.2$ | $\text{mm}^3$ |
| Vascular Density | 5.07 | 0.37 | % |

### Computational Model

Our computational framework is compartmentalised into two models. The first predicts blood flow through the tumour vasculature and the second predicts interstitial fluid flow throughout the cancerous tissue through use of Green’s functions. Our method enables application to whole, large vascular networks (> 2 cm^3^ with > 10^5^ blood vessels), thereby permitting predictions of whole tumour fluid dynamics which incorporate the inherent architectural heterogeneities occurring at the micron-scale.

The intravascular component incorporates the model of Pries et al. [42] to simulate vascular blood flow, where the structural properties of the segmented tumour networks and haemodynamic parameters are used as inputs. Flow or pressure boundary conditions at all terminal nodes in the vascular network are required to predict blood flow throughout the network. These boundary data are very challenging to measure *in vivo*, so we deploy the flow estimation algorithm of Fry et al. [39], to estimate boundary data based on the assumption that the microcirculation is regulated in response to haemodynamics stimuli relating to flow and shear stresses [43]. The scheme estimates unknown boundary conditions by minimising the squared deviation from specified target network wall shear stresses and pressures values derived from independent information about typical network haemodynamic properties. In essence, the algorithm removes the need to define conditions at all boundary nodes, to one where simulation sensitivity is weighted towards the definition of these two target parameters. This enables physiologically realistic blood pressure and flow distributions to be estimated across an entire vascular network and has been applied to breast tumour [44], colorectal tumours [38], cortex [45], glioma [38] and skeletal muscle [46].

The second component to our computational model describes fluid transport through the porous interstitium using a Darcy model, coupled to the vascular flow solution via Starling’s law which describes fluid transport across the endothelium. The vasculature is represented by a discrete set of points sources of flux where the source strengths are defined by the vascular blood flow solution. A similar approach has been applied to simulate O_2_ transport across various tissues [40, 45, 47]. Our approach enables us to explore the effect of vascular architecture heterogeneity on fluid transport within the interstitium for large-scale vascular networks.

The following sections detail the mathematics behind our model along with its computational implementation for large vascular networks. We detail the assignment of model parameters and boundary conditions for application of our model to tumour networks.

#### Vascular Blood Flow

The segmented tumour networks consist of a series of vessel segments connected by nodal junctions or, in the case of boundary nodes, one-segment nodes which form a boundary to the microvascular network (see Fig 1 b). We define a positive flow direction from the start node to end node of each vessel segment. Under the assumption of Poiseuille flow and conserving flow at blood vessel junctions, the relationship between nodal pressures, *p_k_* and the boundary boundary fluxes *Q*_0*i*_ is given by

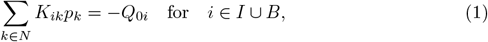

where *N* is the set of all nodes, *I* is the set of all interior nodes and *B* is the set of all boundary nodes with known boundary conditions. For all interior nodes, conservation of flux at vessel junctions dictates that *Q*_0*i*_ = 0, however, if *i* is a known boundary node, *Q*_0*i*_ is the inflow (or outflow if negative).

Following the notation of Fry et al. [39], the matrix *K_ik_* represents network conductance

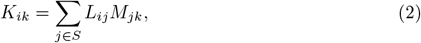

where *S* is the set of all segments,

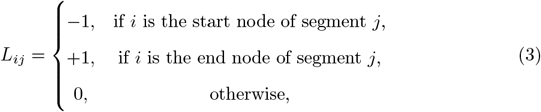

and

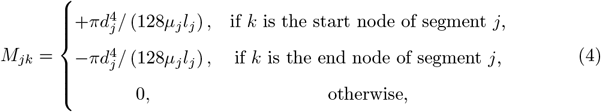

is the matrix of vessel conductances where *l_j_, d_j_* and *μ_j_* denote the length, diameter and effective blood viscosity of segment *j*, respectively.

We apply empirical *in vivo* blood viscosity laws, which prescribe the effective blood viscosity as a function of vessel diameter and haematocrit, to compute *μ_j_* and consequently incorporate non-Newtonian effects in each individual microvessel [48]. Network haematocrit heterogeneity plays an important part in network flow resistance. However, in this study, we set network haematocrit to 0.45 as we do not have sufficient data to parametrise a haematocrit model at this scale. With future availability of appropriate data, the model has the flexibility to incorporate haematocrit heterogeneity [49].

In the absence of measured flow and pressure data at network boundaries, further assumptions are required to obtain a unique solution. The method proposed by Fry et al. [39] sought to solve a constrained optimisation problem, formulated in terms of a Lagrangian objective function defined by

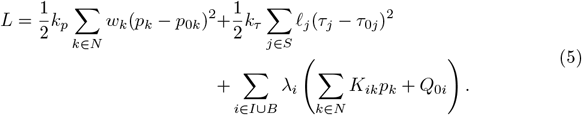

Here, *p*_0*k*_ is the target pressure at node *k, τ_j_* is the wall shear stress in segment *j, τ*_0*j*_ is the corresponding target shear stress, *k_p_* and *k_τ_* are weighting factors associated with the pressure and shear deviations from the target values, *λ*_*i*_ is the Lagrange multiplier associated with node *i* and *w_k_* is the vessel length associated with node *k*. Setting *dL/dp_i_* = 0 and combing with (1) yields a sparse linear system with unknowns *p_k_* and *λ*_*i*_. Assigning a pressure drop to a proportion of boundary nodes forms a well-posed system which can be solved using standard methods [39].

The blood flow estimation model by Fry et al. [39] has been thoroughly tested using mesenteric networks [50] in which blood flow measurements were taken in individual vessels and used to inform parameter estimation [39, 42, 48, 51].

#### Interstitial Fluid Transport

Darcy’s law has been effectively used to describe the passage of fluids [3, 14, 30–33, 52] or solutes [40, 45] through tissues. In this study, we use Darcy’s law to describe the relationship between the volume-averaged IFP, *p*, and interstitial fluid velocity (IFV), **u**, within the porous interstitial domain:

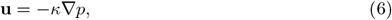

where *κ* is the hydraulic conductivity of the interstitial tissue. Here we assume that interstitial pressure tends towards a constant value, *p*_∞_, in the far-field region,

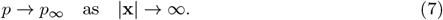

Tumours are leaky due to large pores along a vessel’s lumen, and so the vasculature exhibits a strong fluid and oncotic interaction. Following the approach of Baxter and Jain [3] and subsequent studies [53, 54], we used Starling’s law to describe fluid transport across the endothelium:

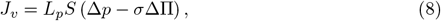

where *J_v_* and *L_p_* is the fluid flux across and the hydraulic conductance of the vessel wall, respectively, *S* is the surface area of the vasculature, *σ* is the oncotic reflection coefficient and, Δ*p* and ΔΠ are the fluid and oncotic pressure gradients between the vasculature and tissue.

The tumour vascular architecture is used to spatially parametrise the locations of a discrete set of sources of flux into the interstitial domain. Assuming these sources both supply or drain the interstitium, conservation of mass yields

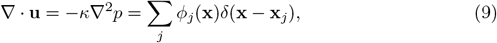

where **x**_*j*_ and *ϕ_j_* are the spatial coordinates and strength, respectively, of point source *j*, and *δ*(**x** − **x**_*j*_) is the delta function. The term *ϕ_j_*(**x**)*δ*(**x** − **x**_*j*_) represents a point source of fluid flux from the vasculature to the surrounding interstitial domain.

Applying the substitution 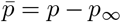, the Green’s function, *G*(**x, x**\*), for the adjoint problem for 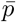 is given by

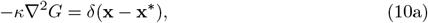

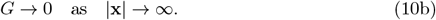

For source a given source, we distribute the delta function over a sphere of finite radius, *r*_0_. This allows the corresponding Green’s function to be described as a radially symmetric function:

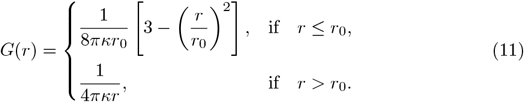

We distribute a total of *N_s_* sources across the vasculature with spacing *λ*_*i*_, and assign a source radii, *r*_0*i*_, equal to the corresponding blood vessel radius (see Fig 1 c). Using Green’s superposition principle for linear operators, the convolution of *G* provides the corresponding pressure solution for source *i* ∈ *N_s_* and so

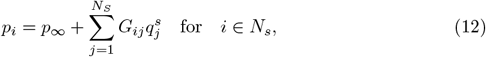

where 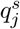 is the vector of source strengths. Here, *G_ij_* is the Green’s function associated with (10) and defined by

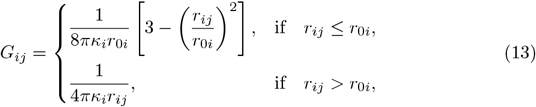

where *r_ij_* is the distance between sources, defined as *r_ij_* = |**x**_*i*_ − **x**_*j*_|, and *κ_i_* is the interstitial hydraulic conductivity at the location of source *i*.

From (6), the Green’s function, *G_ij_*, can be used to calculate the volume-averaged IFV, given by

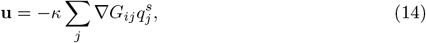

where

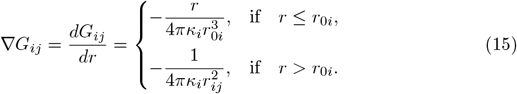

Starling’s law, (8), can be rearranged into the form

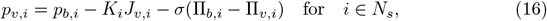

where *p_v,i_* (calculated by (12)) and Π_*v,i*_ are the blood and oncotic plasma pressure at the vessel wall, *p_b,i_* (calculated by (5)) and Π_*b,i*_ are the vascular blood pressure at source i, in the absence of diffusive interstitial fluid transfer, and oncotic fluid pressure, and *J_v,i_* is the rate of fluid flow per unit volume from blood vessel *i* to the interstitium. The intravascular resistance to fluid transport across blood vessel *i*, is defined by

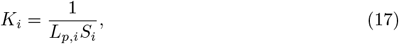

where *S_i_* and *L_p,i_* is the surface area and vascular conductance of vessel *i*, respectively.

Integrating over the volume of vessel *i* and assuming flux at the interface is continuous, flux at the interface is defined by

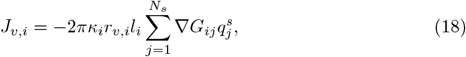

where *r_v,i_* is the vessel radius and *l_i_* is the length of vessel *i*.

Equations (12), (16) and (18) are then combined to give a set of *N_s_* equations to be solved for the source fluxes 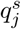,

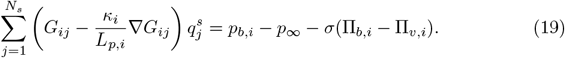

Prescribing parameter values (see Table 2), the resulting solutions for 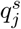 can be solved and used to update vascular pressures, *p_i,b_*, using Starling’s law, (8), in the absence of an oncotic pressure gradient (i.e. ΔΠ = 0), and (18). Here, for iteration *k* + 1, *p_i,b_* is set equal to the IFP at wall of blood vessel *i, p_i,v_*, calculated on iteration *k*. This iterative system is repeated by updating 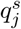 values until tolerances are reached 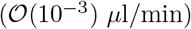, subsequently, tissue IFP and IFV fields can be computed using (12) and (14), respectively (see Fig 1 d).

**Table 2.** Fluid Transport Parameters

| Parameter | Description | Value | Units | Reference |
| --- | --- | --- | --- | --- |
| $p_\infty$ | Far-field interstitial fluid pressure | 0 | mmHg | - |
| $\kappa$ | Interstitial hydraulic conductivity | $1.7 \cdot 10^{-7}$ | $\text{cm}^2 \text{mmHg}^{-1} \text{s}^{-1}$ | [17] |
| $L_p$ | Vascular hydraulic conductance | $2.8 \cdot 10^{-7}$ | $\text{cm mmHg}^{-1} \text{s}^{-1}$ | [11] |
| $\sigma$ | Oncotic reflection coefficient | 0.82 | - | [60] |
| $\Pi_b$ | Oncotic pressure of blood | 20 | mmHg | [61] |
| $\Pi_v$ | Oncotic pressure at the vessel wall | 15 | mmHg | [11, 62] |
| $S/V$ | Ratio of vascular surface area to tumour volume | | | |
| GL261 | | 17.3 | $\text{cm}^{-1}$ | Calculated. |
| LS147T | | 15.3 | $\text{cm}^{-1}$ | Calculated. |

Assigned baseline parameters for the interstitial component of the fluid transport model. Note, $S$ was calculated using the architectural data of the vasculature and $V$ based on computing the convex hull of the tumour.

In effect, our computational framework enables a detailed, quantitative assessment of blood and interstitial flow for tissues, both healthy and pathological, where their entire vascular networks are characterised, by encapsulating the flow of fluid between the vascular and interstitial domains. Similar to a Green’s function model for O_2_ transport [40], our model does not require the imposition of explicit boundary conditions on the outer surface of the tissue domain, with the only unknowns to the system being the strength of the set of fluid sources and sinks. As such, when compared to finite difference or element methods, our approach minimises boundary condition artefacts and saves on computational expense as the solutions to the entire mesh are not required. An outline of the interaction between the biomedical imaging, and vascular and interstitial flow compartments is given in Fig 1 d.

The computational framework was coded using C++ [55] and run on a Apple Mac Pro, with 2 × 3.06 GHz 6-Core Intel Xeon processor and 64 GB of RAM. The system (19) was used solved using a biconjugate gradient method [56] and implemented using the Armadillo sparse linear algebra library [57].

### Implementation of Computational Model on Whole Tumour Microvascular Networks

It remains practically infeasible to measure vascular flows and pressures in individual microvessels *in vivo*, which necessitates a pragmatic approach to boundary condition assignment. Under the assumption that vessels along the tumour surface are connected to peritumoural vessels [44, 58], we developed an optimisation procedure which assigns vascular pressures to tumour surface vessels, based on a target pressure drop, with iterative adjustments to match *in vivo* measurements of mean perfusion from ASL-MRI (see Fig 2 a, b). These *in vivo* data are acquired for the same tumours that were subsequently subjected to OPT analysis. Using this approach, we are able to ensure good agreement between *in silico* predictions and measured perfusion data [38].

**Fig 2.**
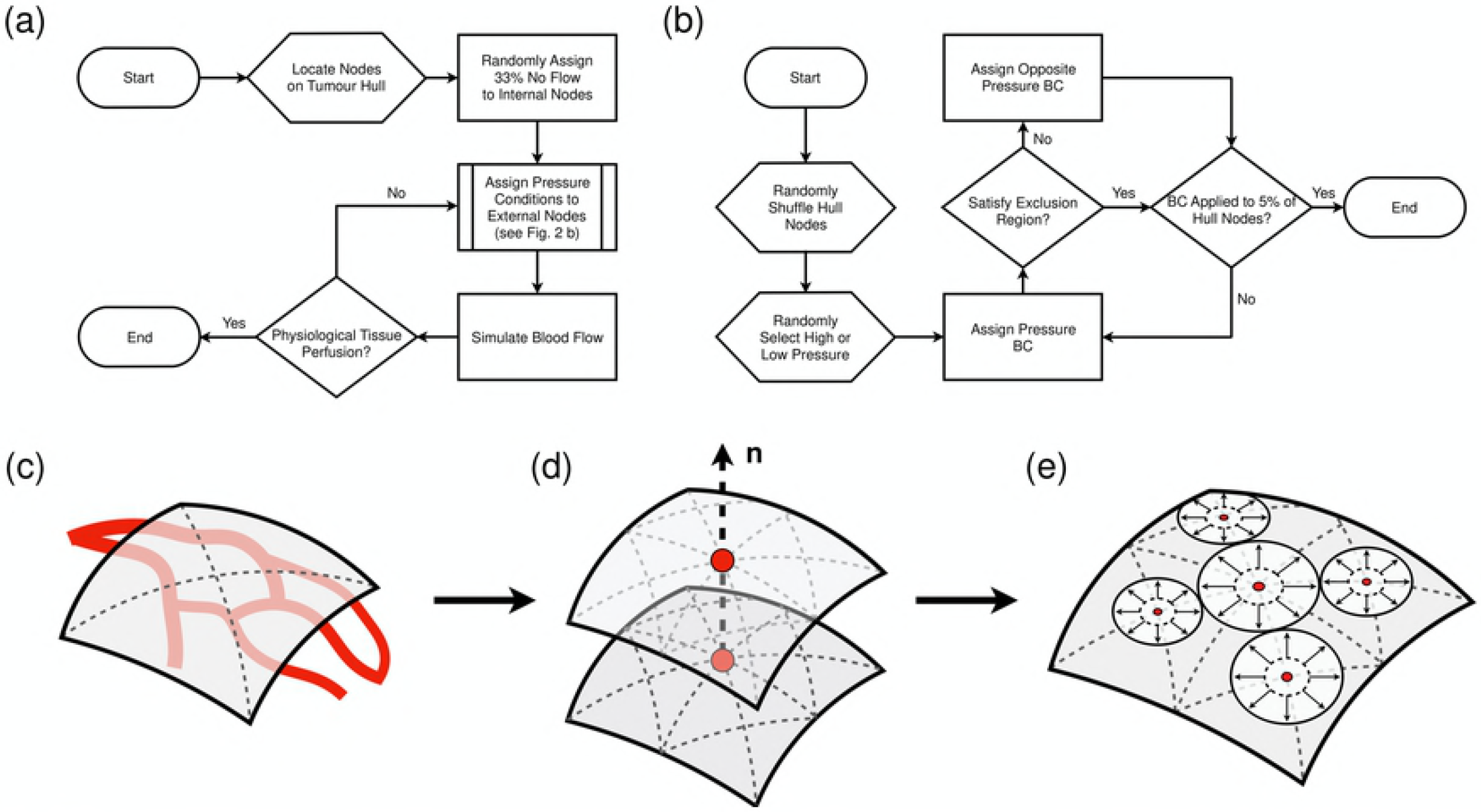
(a, b) The optimisation scheme used to assign boundary conditions to the tumour networks. (a) The process to simulate physiological tissue perfusion. (b) The flowchart for the subroutine “Assign Pressure Conditions” given in (a). (c) Perfusion through a tumour is calculated by generating a convex hull across the surface of the tumour to accurately extract tumour volume. (d) Discretising the hull into a finer mesh and calculating IFP at coupled points across, and normal, to the tumour hull. (e) A sphere packing algorithm is then applied to the points on the tumour surface with inflow averaged across the great circles of each sphere, enabling an approximation of perfusion.

In this study a vascular pressure of 30 or 20 mmHg for the GL261 and 45 or 15 mmHg for the LS147T tumour was randomly assigned to 5% of surface boundary nodes, in order to meet the required tissue perfusion. To ensure randomness, the peritumoural nodes were represented by a list. The elements in the list were rearranged randomly using a uniform random number generator where the system clock was used to seed the random number engine. The nodes located in the top 5% of the list were then randomly assigned a low or high pressure use an equivalent randomised approach.

During preliminary simulations it was found that if high/low pressures were prescribed in close local proximity to each other, unphysiological flows were predicted due to the steep local vascular pressure gradient. In order to prevent this, a subroutine was designed so that values at opposing ends of the pressure spectrum could not be located within a defined vicinity of each other. This “exclusion region”, centred on a given boundary node, was defined as an ellipsoid volume with diameters, along its three principal axes, equal to 5% of the tissue dimensions.

A proportion of boundary nodes are contained within the tumour volume as a consequence of either angiogenesis which form blind-end vessels, or artefacts of OPT. Previous studies approximated the fraction of blind-end vessels in sample MCa–IV carcinomas to be 33% [59]. However, blind-end information is not available for either GL261 or LS147T tumours. Therefore, consistent with previous computational studies [44], blind-ends were randomly applied to 33% of remaining boundary nodes (using the previous randomised approach), with the remaining 62% of boundary nodes left as unknown in the flow optimisation scheme of Fry et al. [39].

Application of our boundary assignment method requires us to accurately compute mean perfusion across the tumours for a comparison against equivalent experimental data gathered *in vivo* using ASL-MRI. This requires an accurate definition of the tumour surface and volume to give an accurate approximation of a tumour’s mass and fluid flow into the tumour volume. For example, an overestimation of the tumour mass, assuming a cuboid tissue volume surrounding the tumour, can drastically underestimate tumour perfusion since, in this case, the tumour shapes are approximately ellipsoidal and perfusion is inversely proportional to the tumour mass. Similarly, an overestimation of flow into the tissue would overestimate tissue perfusion. Next we describe: 1) defining the surface of the tumour; 2) computing the IFV vectors across the tumour surface; and 3) approximating the total tissue perfusion.

1) The hull of a tumour is calculated using the Matlab (MathWorks Inc., Natick, MA) ‘*boundary*’ function applied to all nodes defined during vascular segmentation (see Fig 2 c). The Matlab ‘*fast loop mesh subdivision*’ triangulation algorithm is then applied to further discretise the define tumour surface. 2) To approximate tumour perfusion requires us to define a set of normal vectors to the tumour surface to compute pressure gradients. We identify the centre of the tumour and duplicate the hull, which is then expanded to form a 10 *μ*m gap between the two surfaces (see Fig 2 d). IFP is then calculated across all nodal points on each surface, each paired by a vector normal to the opposing surface. A pressure gradient is then computed along each normal vector and the corresponding velocities are calculated. 3) A sphere packing algorithm is applied to the nodes on the original tumour hull, whereby no sphere overlaps neighbouring spheres (see Fig 2 e). Any inflowing node (defined by the corresponding pressure gradient) is averaged across the great circle of its corresponding sphere and its contribution is summed together to calculate the interstitial component of tissue perfusion. Total tissue perfusion is calculated by summing over the peritumoural vascular inlets and interstitial perfusion values.

Finally, we prescribe baseline parameter values (see Table 2). We assume that the tumours were isolated in subcutaneous tissue in the absence of lymphatics, therefore the far-field pressure, *p*_∞_, was set to 0 mmHg. Due to a lack of experimental data, the vascular conductance and interstitial conductivity, *L_p_* and *κ*, respectively, were given uniform values based on literature (2.8 × 10^−7^ cm mmHg^−1^ s^−1^ and 1.7 × 10^−7^ cm^2^ mmHg^−1^s^−1^, respectively [31, 63]). As the transport of blood plasma proteins is not modelled explicitly in our model, we assume a constant oncotic pressure gradient of 5 mmHg between the vasculature and interstitium.

## Results

In the following section we apply our computational framework to a GL261 orthotopic murine glioma and a LS147T human colorectal carcinoma xenograft to form baseline flow solutions. We then explore sensitivity to source parameters, which include source distribution, source radii and bilateral communication between the vascular and interstitial compartments. We then perform sensitivity analysis to the interstitial parameters (for example, vascular hydraulic conductance and interstitial hydraulic conductivity) on IFP and IFP profiles in the LS147T tumour. We also provide an example of changes occurred by parameter variation in fluid dynamic spatial maps.

### Simulating Interstitial Fluid Pressure in Real-World Cancerous Tissue

Our vascular flow simulations are in good agreement with those in computational [44, 64] and experimental literature (see Table 3 and Fig 3). To test the variability induced by our stochastic boundary condition implementation, the optimisation procedure (detailed in Fig 2 a, b) was repeated for a total of n = 12 for each tumour. Interstitial fluid flow was then simulated for each separate vascular flow solution, providing us with a baseline set of interstitial flow solutions. Blood flow across all simulations exhibited similar spatial distributions, with perfused vessels mainly restricted to the outer rim of the tumours [65]. The mean standard deviation of vascular blood pressures across all simulations was low with values of 0.82 and 1.53 mmHg (with maxima of 4.78 and 13.2 mmHg) for GL261 and LS147T, respectively (see Fig 5 a, e). The elevated standard deviations in vascular pressure were located at the periphery of the tumours, which is to be expected as the high and low vascular pressures were stochastically assigned here. Furthermore, our mean blood velocity and vessel wall shear stresses agree with similar numerical modelling of vascular blood flow in the MDA-MB-231 breast cancer cell line [44].

**Fig 3.**
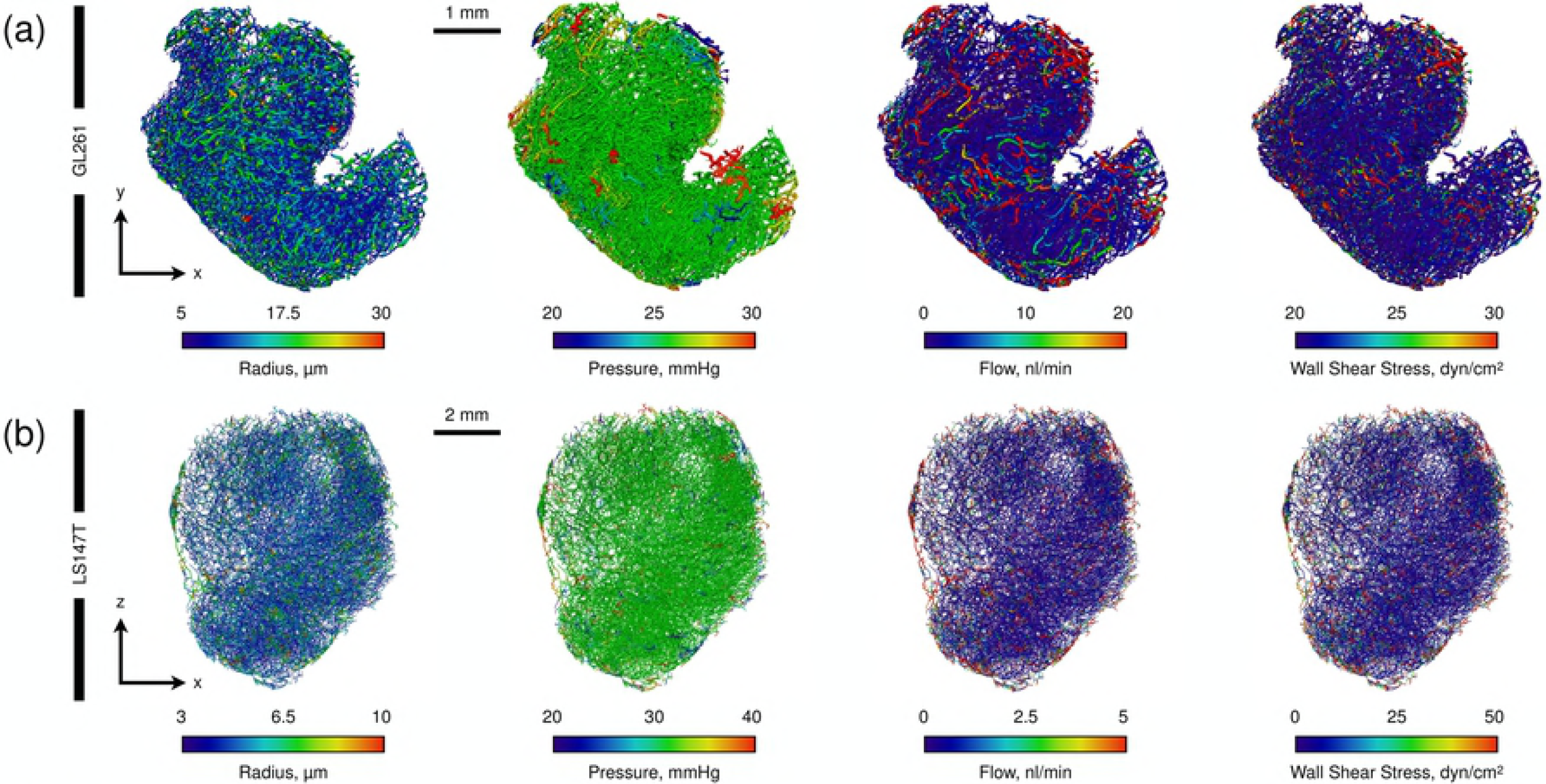
Simulated vascular blood flow in (a) GL261 and (b) LS147T tumours. Distributions are shown for vessel radii, blood pressure, flow and vessel wall shear stress, respectively.

**Table 3.** Simulated Fluid Transport Statistics

| Parameter | GL261 | LS147T | Literature | Units |
| --- | --- | --- | --- | --- |
| Blood Pressure | $25.0 \pm 0.9$ | $30.6 \pm 2.0$ | 10 – 80 [64] | mmHg |
| Blood Flow | $5.1 \pm 43.4$ | $1.1 \pm 4.7$ | 0 – 180 [64] | nl min<sup>-1</sup> |
| Blood Velocity | $0.2 \pm 1.0$ | $0.2 \pm 0.6$ | $1.62 \pm 0.14$ [44] | mm s<sup>-1</sup> |
| Vessel Wall Shear Stress | $5.7 \pm 23.2$ | $11.8 \pm 32.6$ | $23.9 \pm 0.7$ [44] | dyn cm<sup>-1</sup> |
| Tissue Perfusion | $83.7 \pm 22.3$ | $5.4 \pm 0.3$ | $110 \pm 70^{\dagger}$, $19 \pm 8^{\ddagger}$ [38] | ml min<sup>-1</sup> 100 g<sup>-1</sup> |
| Interstitial Fluid Pressure (IFP) | $20.4 \pm 2.1$ | $25.3 \pm 1.6$ | $13.5 \pm 11.3$ [62]<sup>‡</sup> | mmHg |
| Interstitial Fluid Velocity (IFV) | $4.8 \pm 0.5$ | $0.19 \pm 0.09$ | 2.3 [25] | $\mu\text{m s}^{-1}$ |

Intra- and extravascular baseline fluid transport statistics for the GL261 and LS147T tumours (mean ± standard deviation) across all simulations and compared against literature values. <sup>†</sup>: Data gathered from GL261 tumours. <sup>‡</sup>: Data gathered from LS147T tumours.

Tissue perfusion (calculated as 83.7 ± 22.3 and 5.4 ± 0.3 ml/min/100g for the GL261 and LS147T tumours, respectively) was further validated by *in vivo* ASL-MRI measurements of at 110 ± 70 and 19 ± 8 ml/min/100g for GL261 and LS147T, respectively. This implies that imposing physiologically realistic pressure boundary conditions generates physiologically realistic perfusion, and consequently accurate drug delivery solutions [38].

No literature IFP values were available for GL261 cell lines, however, our IFP was slightly elevated compared to that previously measured in LS147T *in vivo* (13.5 ± 11 . 3 mmHg [62]). Considering the range of IFP both here and *in vivo* [62] and the good accordance with *in vivo* perfusion, our results provided us with the confidence that our simulations can produce physiological IFP predictions (see Fig 4).

**Fig 4.**
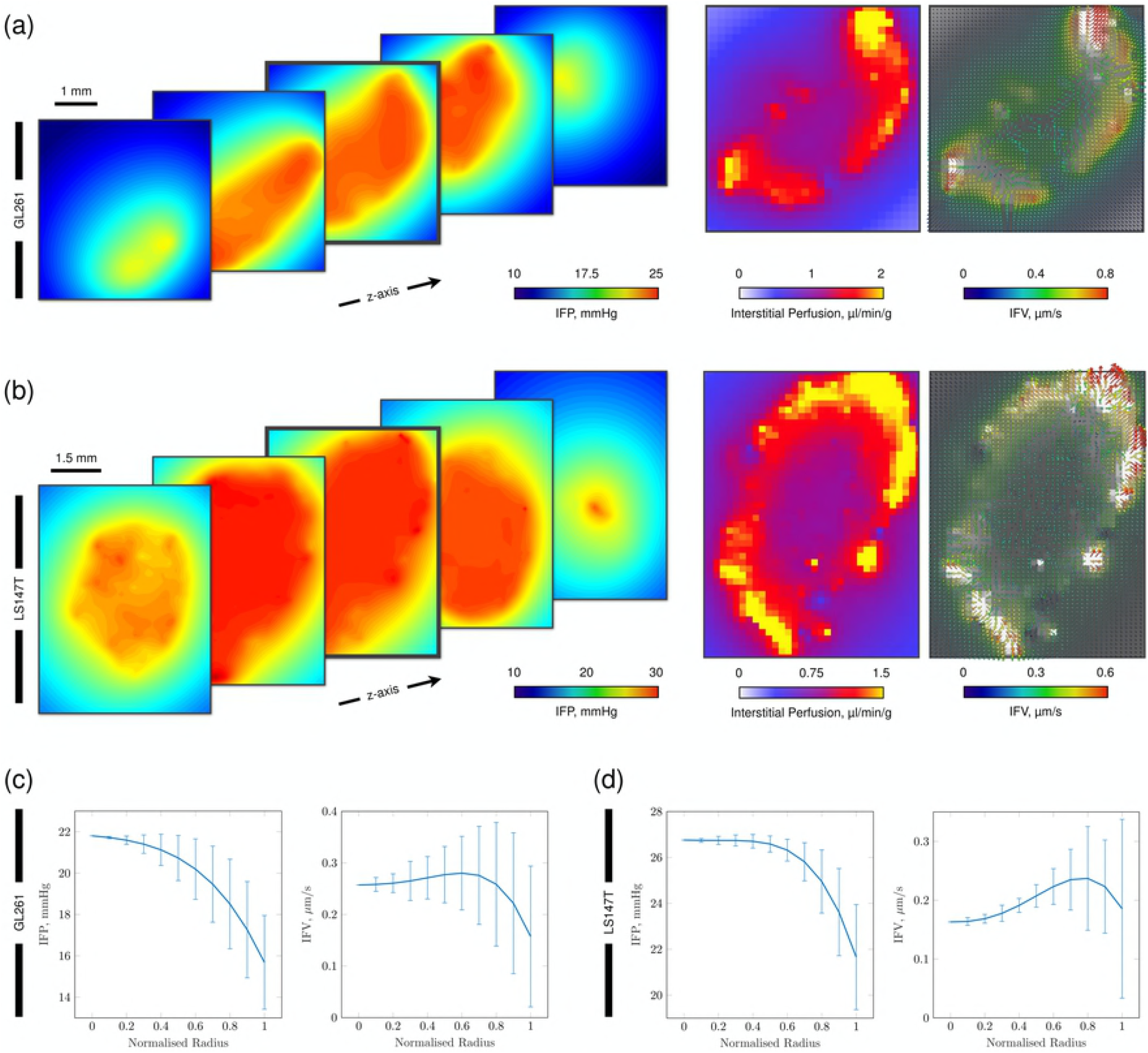
Simulated fluid transport through the interstitium in GL261 and LS147T tumours. (a, b) (Left) Predicted interstitial fluid pressure (IFP) fields for {X, Y}-planes through the tumours, emulating the traditionally high pressure in the tumour core but with predicted spatial heterogeneities. (Middle) Simulated interstitial perfusion maps discretised into ~ 140 *μ*m^2^ pixels. Results replicate the traditionally elevated perfusion existing at the periphery of the tumours. (Right) Interstitial fluid velocity (IFV, overlaid onto greyscale image of interstitial perfusion) predictions depicting spatial interstitial flow heterogeneities across the entire tumours. Note, perfusion and interstitial fluid velocity maps are shown for the central slice in the interstitial fluid pressure graphics. (c, d) Fitted curves with error bars indicating standard deviation for (left) IFP and (right) IFV in (c) GL261 and (d) LS147T, plotted against normalised radius, corresponding to the simulations shown in (a, b).

Examining the tumour radial IFP profiles, the LS147T network exhibited similar configurations as observed both experimentally in LS147T [17], in other cell lines [3, 24, 25], and in computational studies [31, 32, 36, 63], with elevated IFP at the tumour core (see Fig 4 b, d). In addition, the LS147T network displayed a typical IFV profile radially, with an increasing IFV range towards the tumour surface due to the steeper pressure gradients at the periphery of the tumour. This indicated that bulk fluid filtration occurs at the high flowing vasculature located at the tumour extremity in this network (*ρ* = 0.41, *p* < 0.001, where *ρ* and *p* are the Pearson’s correlation coefficient and its corresponding *p*–value between tumour radius and vascular flow, respectively).

In comparison, the GL261 network also exhibited a traditional, yet steeper, IFP profile with a wider variance throughout the tumour (see Fig 4 c). The IFV profile exhibited a typical profile increasing from the centre of the tumour to the periphery. However, its IFV peak was reached at ~ 80% of the tumour radius, with a substantial decline in the latter 20% (see Fig 4 c).

It was observed that GL261 and LS147T have distinct differences in their vascular architecture (see Fig 3 and Table 1). This may indicate that the inherent vascular heterogeneity across tumour cell lines [20] directly impacts the IFP and IFV distributions, creating an unorthodox interstitial flow profile. High IFP has been associated with low vascular density in A-07-GFP tumours [66], however, further work is required to confirm our hypothesis for GL261 and LS147T tumours.

### Sensitivity of Tumour Interstitial Fluid Transport to Model Parameters

Sensitivity analysis allows us to understand the impact of parameter variance on tumour perfusion predictions. In this study, we performed sensitivity analysis to our model’s underlying parameters. For convenience we have split these parameters into two groups. The first we define as source parameters, which include the vascular blood pressure, *p_b,i_*, the spacing between each source along a given vessel, *λ*_*i*_, and the source radius, *r*_0,*i*_ for segment *i* (see Fig 1 c). The second group we call the interstitial parameters, which include the far-field IFP, *p*_∞_, the oncotic reflection coefficient, *σ*, the hydraulic conductance of a vessel wall, *L_p_*, and the interstitial hydraulic conductivity, *κ*. In the following, if sensitivity analysis to an interstitial parameter was not being performed, it was set to the corresponding value in Table 2.

#### Source Parameters

Vascular pressure depends not only on the vascular pressure at the boundary nodes and vascular network geometry but also on the permeability of the tumour vessels and hydraulic conductivity of the interstitial tissue, as a consequence of flow communication between the vascular and interstitial domains. To incorporate this relationship, we couple the vascular and interstitial models using an iterative scheme in which vascular blood pressure distributions, *p_b,i_* for *i* ∈ *N_s_*, were updated on each iteration by incorporating the loss of fluid within a vessel due to fluid flux across the vessel wall into the interstitium (see Fig 1 d). We found that due to the small volume of fluid leaving the vessels, vascular pressure was not significantly corrected on each iteration (see Fig 5 b, f). In the case of the GL261 network, the algorithm had converged after two iterations (see Fig 5 b). In comparison, the LS147T network had not converged after six iterations but the mean error was 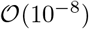 mmHg (see Fig 5 f). This may indicate that a tumours inherent vascular heterogeneity and corresponding interstitial parameters alter the scale of these flow errors.

**Fig 5.**
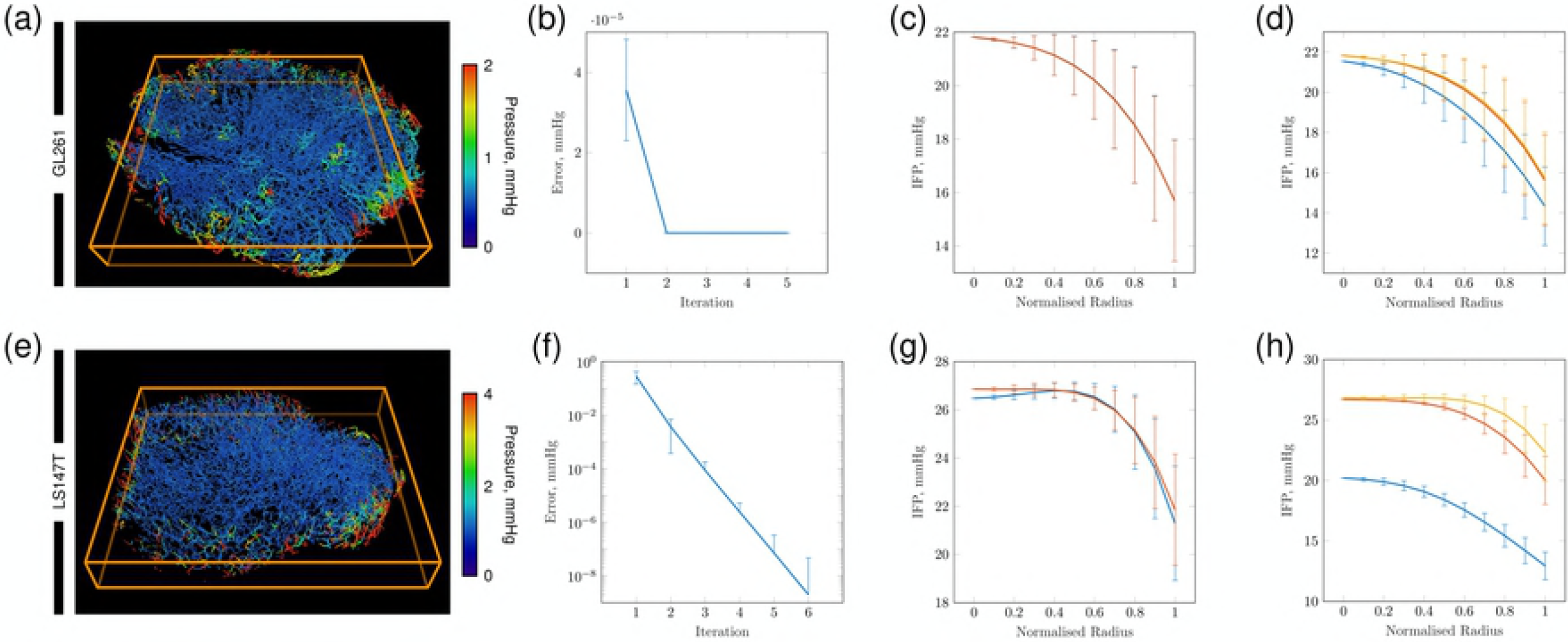
Sensitivity of the computational framework. (a, e) Cross-sectional slice from the core of the (top) GL261 and (bottom) LS147T tumours, showing standard deviation of intravascular pressures (mmHg) across all 12 simulations. (b, f) the mean segment pressure error between consecutive iterations for the (top) GL261 (top) and (bottom) LS147T networks. (c, g) IFP distribution from the centre of the tumour to its periphery for a maximum spacing of (blue) 10 and 25 *μ*m and (orange) 50 and 100 *μ*m for GL261 and LS147T, respectively. (d, h) IFP distributions for the source radii set to the minimum vessel radius multiplied by a factor of (blue) 10^−1^, (red) 10^0^ and (yellow) 10^1^. Note, error bars correspond to standard deviation.

Next we varied the maximum distance, *λ*_*max*_, between sources distributed along the vasculature. Values of 10 and 25 *μ*m, and 50 and 100 *μ*m were chosen for the GL261 and LS147T networks, respectively, due to differences in mean branching vessels lengths observed in each tumour (see Table 1). The GL261 network exhibited minimal sensitivity in IFP between the two maximal source lengths (see Fig 5 c). Comparatively, the LS147T experienced greater variability (of less than 1 mmHg) in mean IFP, between the two values of *λ*, at the core and periphery of the tumour (see Fig 5 g). We hypothesise that sensitivity to *λ* is related to the vascular density of each tumour type. The GL261 network is an order of magnitude greater in vascular density compared to the LS147T tumour (see Table 1). An elevated vascular density results in an increased density of fluid sources, and so any reduction in sources may have a minimal effect to IFP due to levels being maintained by the local proximity of neighbouring sources.

We finally sought to understand how the source radii, *r*_0,*i*_ for *i* ∈ *N_s_*, affects flow in the interstitial domain. We initially set the source radii to a constant value equal to the minimum vessel radius in a given tumour network. Three cases were then explored where *r*_0,*i*_ was multiplied by a factor of 10^−1^, 10^0^ or 10^1^. Our results show that decreasing the value of the source radii decreases mean IFP, with greater sensitivity exhibited again by the LS147T network (see Fig 5 d, h). This indicates that greater care is needed compared to the assignment of other source parameters, to ensure the physiological accuracy of the results. Here, we set the value of *r*_0,*i*_ equal to the radius of its corresponding vessel in order to provide a heterogeneous distribution of radii in which vessels with a larger radius are able to influence the IFP to a greater capacitys than compared to relatively smaller vessels.

#### Interstitial Parameters

Sensitivity to the interstitial parameters *p*_∞_, *σ, L_p_*, and *κ* were explored using the LS147T network. This network was chosen due to the scale of simulations to be performed and due to the greater sensitivity exhibited by modifying the source parameters. First of all, we investigated the deviation in the far-field interstitial pressure, *p*_∞_. Ideally, the far-field pressure is set to the mean IFP of the adjoining healthy tissue, however these data are frequently unavailable. We varied the far-field IFP from 3 to 18 mmHg in increments of 3 mmHg. Alteration of *p*_∞_ did not significantly modify tumour IFP, with the majority of variation occurring at the periphery of the tumour (see Fig 6 a). As a consequence, mean IFV across the tumour decreased with increasing *p*_∞_ (see Fig 6 e).

**Fig 6.**
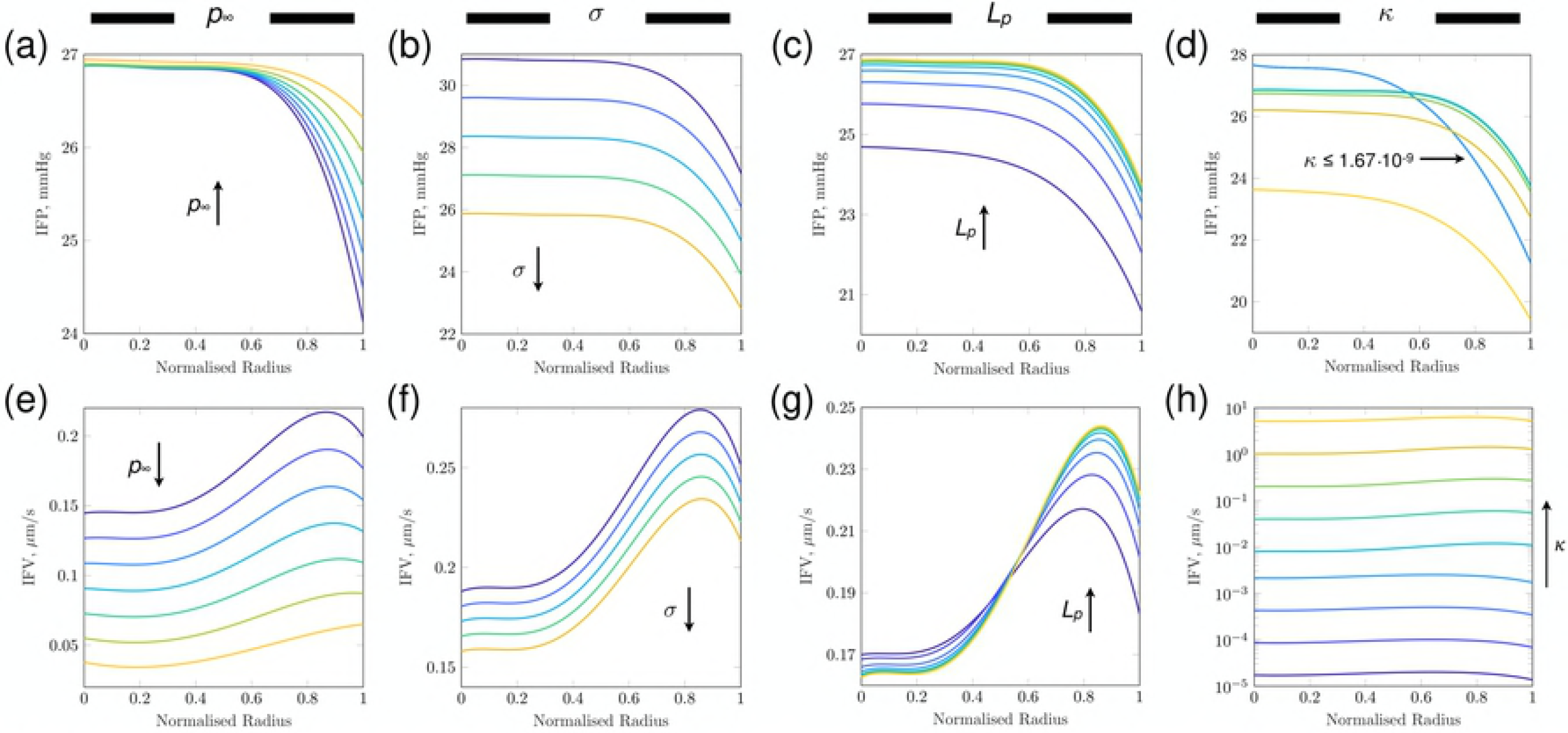
Data-fitted curves of IFP (top) and IFV (bottom) for modulation of interstitial model parameters: (a, e) *p*_∞_ (mmHg), (b, f) *σ*, (c, g) *L_p_*(cm/ mmHgs) and (d, h) *κ* (cm^2^ / mmHgs). Arrows indicate increasing parameter values, with exception of (d) in which a range of *κ* (cm^2^ / mmHgs) is indicated. IFP profile gradients across LS_2_ sensitivity analysis of (a) far-field pressure, *p*_∞_, (b) oncotic reflection coefficient, *σ*, (c) vascular hydraulic conductivity, *L_p_*, and (d) interstitial hydraulic conductivity, *κ* (cm^2^ / mmHgs). Arrows indicate increasing values of the given parameter and colours in each column indicate the equivalent simulation.

Next we investigated the oncotic reflection coefficient, *σ*. Physiologically, the coefficient *σ* varies between 0 and 1, indicating the likelihood that a molecule approaching a pore in the vessel lumen will be reflected back and thereby retained in the vascular compartment. As such, we ranged *σ* from 0 to 1 with increments of 0.25. Modification of *σ* resulted in a network-wide shift in tumour IFP magnitude, thereby preserving the IFP gradient (see Fig 6 b). This is an intuitive result since we held the oncotic pressure gradient constant across the entire vasculature, therefore any increase in *σ* resulted in systematic reduction in both IFP and IFV (see Fig 6 b, f).

Vascular hydraulic conductance, *L_p_*, defines the leakiness of the lumen to the transport of blood plasma. Our sensitivity analysis varied *L_p_* from 1.33 × 10^−8^ to 3.41 × 10^−6^ cm/mmHg s, encompassing values provided in literature for normal and tumour tissue [11]. Similarly to *σ*, a reduction in *L_p_* resulted in decreasing IFP across the tumour but with contrasting regional gradients (see Fig 6 c). Our analysis showed that with decreasing *L_p_*, IFP tended towards the assigned far-field pressure due to decreasing fluid filtration from the vasculature, and so in its limit, the IFP would uniformly be equal to *p*_∞_. This is due to decreasing fluid filtration, indicating that the model produces a physiologically viable response. In comparison, a reduction in Lp induced minimal changes to IFV in the core of the tumour but resulted in decreases towards the periphery (see Fig 6 g).

Variations in the interstitial conductivity, *κ*, ranged from 1.33 × 10^−11^ to 1.04 × 10^−6^ cm^2^ /mmHg s. In this case, we established that sensitivity to this parameter are non-trivial to elucidate. An inflexion point was observed for values 1.67 × 10^−9^ ≤ *κ* ≤ 8.33 × 10^−9^ cm^2^ /mmHgs in which the spatial IFP distribution switched from one configuration to another (see Fig 6 d). For *κ* ≤ 1.67 × 10^−9^ cm^2^ / mmHg s, IFP solutions displayed near identical configurations (mean IFP of 25.8 ± 2.4 mmHg for all four cases) in which IFP displayed an increased gradient compared to greater values of *κ* along with decreasing IFV (see Fig 6 h). In comparison, *κ* ≥ 8.33 × 10^−9^ cm^2^ / mmHgs displayed similar IFP profiles with mean IFP decreasing with increasing *κ* (26.2 ± 1.5 to 22.5 ± 1.6 mmHg).

#### Mapping the Effects of Vascular Normalisation

As an example of how our computational framework can be used to predict the effectiveness of cancer therapies, we performed a feasibility study of normalising the LS147T vasculature, by modifying the network vascular hydraulic conductances and oncotic reflection coefficients. We crudely simulated vascular normalisation using a linear function where vascular pores sizes (represented through a combination of changes in *L_p_* and *σ*) in the tumour core are typical of tumour vasculature, and gradually returned to physiological values at the tumour periphery (from 2.18 × 10^−7^ to 0.44 × 10^−7^ cm mmHg^−1^ s^−1^ for *L_p_* and from 0.82 to 0.91 for *σ*).

Simulations predicted a steeper IFP gradient across the tumour. This is contrary to previous studies [67, 68], where IFP was found to reduce when tumour vasculature was normalised. However, these studies normalised the vasculature uniformly across the tumour, and our equivalent sensitivity analysis for uniform changes in *L_p_* observed a similar IFP response (see Fig 6 c).

In response to changes in IFP, our model predicted elevated tumour interstitial fluid speed (IFS - see Fig 7 a), a similar result to [68] who observed that advection is dominant at small pore sizes in two-dimensions. Steeper IFP gradients also increased perfusion and outward facing IFV streamlines (see Figs 4 b and 7 b).

Considering our predicted IFP responses for uniform and radially varying vascular normalisation, we hypothesise that for vascular normalisation therapy to effectively reduce to IFP, blood vessels across the tumour need to be normalised. Further, that if vascular normalisation increases macromolecular drug penetration via the vasculature to the core, outward interstitial advective currents may inhibit the effectiveness of such therapies once delivered to the interstitium. However, these effects need to be explored using a comprehensive model to detail the delivery of normalisation therapies, and consequently macromolecular drug delivery. In addition, the effect of normalisation is likely to be dependent on tumour architectural properties, such as vascular density and interstitial hydraulic conductivity, and so requires study across a range of tumours with spatially varying conductivities.

**Fig 7.**
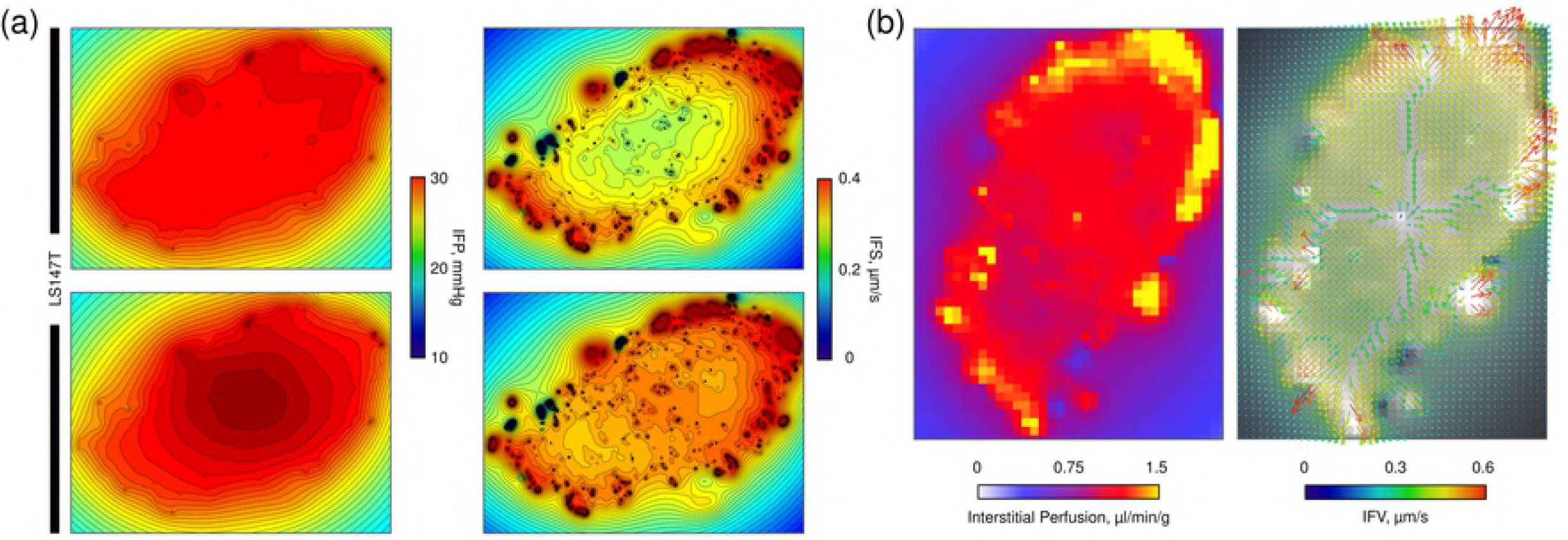
Normalisation of vascular hydraulic conductance and oncotic reflection coefficient to physiological values in LS147T. (a) Planar contour plots of IFP and interstitial fluid speed (IFS) for the baseline (top) and normalised (bottom) predictions. (b) Predictions of normalised interstitial fluid spatial maps (left) and IFV (right - with greyscale perfusion underlaid) where pixels are ~ 140 *μ*m^2^. For comparison, equivalent {X, Y}-planes for baseline simulations can be viewed in Fig 4 c.

## Discussion

Elevated interstitial fluid pressure is frequently associated with solid tumours, where a conventional profile exhibits a uniformly high IFP in the core of a tumour which decreases rapidly towards the levels of physiological tissue at its periphery [3, 14, 52]. This atypical characteristic forms a barrier to transvascular fluid and drug delivery, thereby diminishing therapeutic efficacy of anti-cancer treatments.

The passage of fluid through the interstitium is influenced by both hydrostatic and oncotic pressures in blood vessels and therefore by the heterogeneic architecture of tumour microvessels. However, the procurement of detailed fluid flow data *in vivo* across whole tumour networks is currently unfeasible using conventional imaging and experimental techniques. However, recent advances in *ex vivo* high-resolution optical imaging techniques [37] allow whole three-dimensional tissue architectures to be extracted and reconstructed to act as inputs for detailed *in silico* modelling of fluid transport.

No study has explicitly modelled vascular and interstitial fluid flow using discrete, three-dimensional structural data from images of whole-tumour, real-world vasculature [6]. However, in a recent study, we presented our novel REANIMATE platform which extracts three-dimensional, whole tumour vasculature *ex vivo* from optically cleared tissue, which is then used to parameterise an *in silico* model of fluid transport guided by in vivo imaging data [38]. This has enabled us to perform quantitative, realistic predictions of fluid and drug delivery to tumours which has led to novel insights into a tumour’s inherent physical resistance to anti-cancer therapies [38].

In this study, we presented the computational framework used to simulate fluid transport in REANIMATE. We detail its derivation and application to whole tumour architectural datasets and show how our model allows highly-detailed predictions of fluid flow within the tumour microenvironment by incorporating explicit tumour vasculature. Our model allows flow heterogeneities to be quantified in a computationally efficient manner, when compared to finite-difference and element methods [40], which enables cancer therapies, such as normalisation, to be effectively studied through modification of its parameters.

We initially apply our framework to an orthotopic murine glioma and a human colorectal carcinoma xenografts from the GL261 and LS147T cell-lines, respectively, to present realistic, baseline simulations of the tumour microenvironment. We then performed sensitivity analysis to the underlying model parameters. The first are the source parameters, which are specific to our model, and include source of flux distribution and size. Secondly, the interstitial parameters, such as vascular conductance and interstitial conductivity, which are frequently represented in literature due to the prominent use of Starling’s law. In the second case, we perform analysis of how variation of these parameters modifies the IFP and IFV in an LS147T dataset. We finally present a proof-of-concept detailing how our model can be used to provide three-dimensional, spatial maps of tumour properties, such as IFP, IFS, IFV and perfusion, when investigating the effects of vascular normalisation.

Our computational framework is based upon a Poiseuille model for vascular blood flow [39] which is coupled to a steady-state Green’s function solution to interstitial fluid flow. Here, tumour vasculature segmented *ex vivo* is represented by a discrete set of sources of fluid flux for bi-directional transport between the interstitium. Previous models either assume a homogeneous vascular network [3, 11, 14, 52], incorporate a computer-generated synthetic tumour network [28–34], or incorporate boundary conditions and spatial variations in tissue permeability to artificially represent vascular heterogeneity [35, 36]. However, vascular averaging methods do not fully encapsulating the intrinsic, local interactions between neighbouring blood vessels which contribute to global interstitial flow and synthetic networks are difficult to validate against real tumour architecture. Here we use vascular architecture from real, whole tumour networks, and through use of Green’s function methods, our model significantly reduces the computational size of the computational problem, allowing vessel-vessel interactions to be modelled at the micron-scale. Thus, we provide the means to perform *in silica* studies to hypothesis test the impact of vascular heterogeneity on the tumour microenvironment with relative ease.

Our simulations were performed on *ex vivo* structural imaging data from a GL261 and a LS147T tumour. As no *in vivo* flow or pressure data were available for the numerous boundary vessels, we developed a procedure whereby simulated data was optimised based on *in vivo* tissue perfusion data gathered using ASL-MRI [38]. This approach has produced solutions which are highly consistent with experimental measurements in the same tumours [38]. In this study, baseline vascular flow solutions across all tumour simulations are in good agreement with the perfusion data, alongside mean flows [64], velocities and vessel wall shear stresses [44], and fluid pressure in the interstitium [3, 17, 24, 25]. This provided validation that our model produces physiologically realistic results, providing a platform to investigate the tumour microenvironment.

We performed sensitivity analysis to the source parameters, such as updating the vascular flow solution, source distribution and source radii, to understand their influence on the flow communication between the vascular and interstitial domains. Our results exhibited a minimal sensitivity to IFP distributions by varying these parameters, with the exception of the assignment of source radii. Here, we hypothesise that greater care is required for spatially sparse tumours with a low vascular density, in order to ensure physiologically accurate simulations. Finally we investigate the interstitial parameters using the LS147T dataset. Raising the far-field interstitial pressure did not significantly alter the IFP distribution across the tumour, with increased IFP occurring at the tumour periphery, compared to baseline. Variation in the oncotic pressure contribution, by modifying the oncotic reflection coefficient, only affected the magnitude of the interstitial pressure, with minor changes to the IFP and IFV profiles, similar to those reported in previous studies [3]. Increasing *L_p_* and *κ*, raised the gradient of the interstitial pressure profile, thereby increasing fluid transport through the interstitium. However, our IFP distributions did not reach 0 mmHg at the periphery of the tumours as in previous studies [3, 14, 67]. We hypothesise that this is due to not applying a fixed pressure at the tumour boundary in our model, which results in a smoother transition of IFP to the surrounding tissue, similar to previous computational modelling of tumour vascular heterogeneity [35, 36].

We finally performed a feasibility study detailing how our framework can be used to investigate cancer therapies, such as vascular normalisation. We modified vascular hydraulic conductances and oncotic reflection coefficients in a linear radially varying fashion in the LS147T network. Values ranged from those associated with tumours at its core, to physiological tissue levels at its periphery. Our model predicted that IFP at the core was slightly elevated, and that a pressure gradient is formed radially across the tumour core which causes raised, outwardly directed interstitial flow. Consequently, we predict that vascular normalisation therapies need to uniformly normalise a tumour to lower tumour IFP, and that ineffective normalisation could reduce the efficacy of drug therapies. These results are preliminary, and require further analysis and application to further LS147T (and other cell-line) datasets. However, they indicate that modifying, or normalising, the parameters in our computational framework is a simple and effective way of investigating the fluid dynamic properties, such as interstitial pressures and velocities, associated with an individual tumour. Moreover, our framework predicts spatial maps of flow across three-dimensional cancerous tissue, enabling flow heterogeneities induced by realistic vascular networks to be quantified for subsequent *in silico* investigations into anti-cancer therapies.

Our results and proposed computational framework offer significant scope for future expansion. Current limitations include, for example, our stochastic approach to simulating vascular blood flow which was implemented due to the in-availability of experimental data. Recent *in vivo* methods provide a step forward in approximating conditions by discerning global tumour pressure gradients through observations of tissue fluid velocity [26], which could lead to greater accuracy when assigning boundary conditions specific to a tumour. Similarly, parameter values such as the vascular conductance and interstitial hydraulic conductivity were assigned using previous literature values since these tissue-specific measurements can be challenging to procure through experimentation. For example, interstitial conductivity values across normal tissue have been reported to span four orders of magnitude [17]. Future work could seek to predict spatially heterogeneous maps of interstitial hydraulic conductivity using REANIMATE and new experimental data [26].

There are also opportunities to expand the computational model to incorporate more complex biological phenomena. For example, our model does not currently incorporate tumour compression of vessels due to increasing shear stresses within the tumour, a critical property of understanding angiogenic vasculature [29, 69]. In addition, our model does not explicitly incorporate necrotic regions within cancerous tissues. Yet necrotic regions can influence interstitial fluid delivery [3, 14] and tumour macromolecular transport [52], and so are vital to incorporate if our computational framework.

We expect to find a wide utility for REANIMATE in a range of disease areas, particularly given the current interest in optical clearing methods and their widespread use in biomedical research. REANIMATE is novel and timely and will find extensive use for hypothesis testing, to enable tumour biology and drug delivery to be better understood, which in turn may enable the next generation of cancer therapies.

## Acknowledgements

The authors also acknowledge support received from EPSRC (EP/L504889/1), Rosetrees Trust/Stoneygate Trust (M135-F1 and M601) and the Wellcome Trust (WT100247MA).

## Author Contributions

P.W.S. and R.J.S. developed the mathematical model for fluid transport across tissue and developed the main concepts. P.W.S. developed the software to perform mathematical and computational analysis, analysed and interpreted the results, and wrote and edited the paper. A. d’E designed and performed optical imaging experiments. S.W-S. supervised the design of optical imaging experiments and developed software for segmenting optical imaging data. R.J.S. supervised the design of computational simulations. R.J.S. and S.W-S. co-led the project, secured funding, and contributed to the writing and editing of the manuscript.

## References

1. Andrew I. Minchinton and Ian F. Tannock. Drug penetration in solid tumours. Nature Reviews Cancer, 6(8):583–592, aug 2006. ISSN 1474-175X. doi: 10.1038/nrc1893.

2. Rakesh K. Jain. Determinants of Tumor Blood Flow: A Review. Cancer Research, 48(10):2641–2658, 1988. ISSN 15387445. doi: 10.1146/annurev.bioeng.1.1.241.

3. Laurence T Baxter and Rakesh K Jain. Transport of fluid and macromolecules in tumors. I. Role of interstitial pressure and convection. Microvascular Research, 37 (1):77–104, jan 1989. ISSN 10959319. doi: 10.1016/0026-2862(89)90074-5.

4. R K Jain. Barriers to drug delivery in solid tumors. Scientific American, 271(1): 58–65, jul 1994. ISSN 0036-8733.

5. Seong Hoon Jang, M Guillaume Wientjes, Dan Lu, and Jessie L S Au. Drug delivery and transport to solid tumors. Pharmaceutical research, 20(9):1337–50, sep 2003. ISSN 0724-8741.

6. Heiko Rieger and Michael Welter. Integrative models of vascular remodeling during tumor growth, 2015. ISSN 1939005X.

7. Mark W. Dewhirst and Timothy W. Secomb. Transport of drugs from blood vessels to tumour tissue, nov 2017. ISSN 14741768.

8. Petros Koumoutsakos, Igor Pivkin, and Florian Milde. The Fluid Mechanics of Cancer and Its Therapy. Annual Review of Fluid Mechanics, 45(1):325–355, jan 2013. ISSN 0066-4189. doi: 10.1146/annurev-fluid-120710-101102.

9. S. Goel, D. G. Duda, L. Xu, L. L. Munn, Y. Boucher, D. Fukumura, and R. K. Jain. Normalization of the Vasculature for Treatment of Cancer and Other Diseases. Physiological Reviews, 91(3):1071–1121, jul 2011. ISSN 0031-9333. doi: 10.1152/physrev.00038.2010.

10. Ajit S. Narang and Sailesh Varia. Role of tumor vascular architecture in drug delivery. Advanced Drug Delivery Reviews, 63(8):640–658, jul 2011. ISSN 0169409X. doi: 10.1016/j.addr.2011.04.002.

11. Rakesh K Jain. Transport of molecules across tumor vasculature. Cancer and Metastasis Reviews, 6(4), 1987. ISSN 0167-7659.

12. Rakesh K. Jain. Normalizing Tumor Microenvironment to Treat Cancer: Bench to Bedside to Biomarkers. Journal of Clinical Oncology, 31(17), jun 2013. doi: 10.1200/JCO.2012.46.3653.

13. Michael J Mitchell, Rakesh K Jain, and Robert Langer. Engineering and physical sciences in oncology: Challenges and opportunities, 2017. ISSN 14741768.

14. Rakesh K. Jain and Laurence T. Baxter. Mechanisms of heterogeneous distribution of monoclonal antibodies and other macromolecules in tumors: significance of elevated interstitial pressure. Cancer Research, 48:7022–32, dec 1988. ISSN 0008-5472.

15. Rakesh K Jain. Antiangiogenesis Strategies Revisited: From Starving Tumors to Alleviating Hypoxia. Cancer Cell, 26(5):605–622, 2014. ISSN 18783686. doi: 10.1016/j.ccell.2014.10.006.

16. Carl Henrik Heldin, Kristofer Rubin, Kristian Pietras, and Arne Östman. High interstitial fluid pressure - An obstacle in cancer therapy, oct 2004. ISSN 1474175X.

17. Y. Boucher, C. Brekken, P. A. Netti, L. T. Baxter, and R. K. Jain. Intratumoral infusion of fluid: estimation of hydraulic conductivity and implications for the delivery of therapeutic agents. British Journal of Cancer, 78(11):1442, 1998.

18. Dai Fukumura, Jonas Kloepper, Zohreh Amoozgar, Dan G. Duda, and Rakesh K. Jain. Enhancing cancer immunotherapy using antiangiogenics: opportunities and challenges. Nature Reviews Clinical Oncology, mar 2018. ISSN 1759-4774. doi: 10.1038/nrclinonc.2018.29.

19. Kristian Pietras, Kristofer Rubin, Tobias Sjöblom, Elisabeth Buchdunger, Mats Sjöquist, Carl Henrik Heldin, and Arne Östman. Inhibition of PDGF receptor signaling in tumor stroma enhances antitumor effect of chemotherapy. Cancer Research, 62(19):5476–5484, 2002. ISSN 00085472. doi: 10.1016/j.cell.2009.02.025.

20. A A Folarin, M A Konerding, J Timonen, S Nagl, and R B Pedley. Three-dimensional analysis of tumour vascular corrosion casts using stereoimaging and micro-computed tomography. Microvascular research, 80(1): 89–98, 2010. ISSN 1095-9319. doi: 10.1016/j.mvr.2010.03.007.

21. Yves Boucher, Laurence T. Baxter, and Rakesh K. Jain. Interstitial Pressure Gradients in Tissue-isolated and Subcutaneous Tumors: Implications for Therapy. Cancer Research, 50(15):4478–4484, 1990. ISSN 15387445. doi: 10.16373/j.cnki.ahr.150049.

22. R Gutmann, M Leunig, J Feyh, A E Goetz, K Messmer, E Kastenbauer, and R K Jain. Interstitial hypertension in head and neck tumors in patients: correlation with tumor size. Cancer research, 52(7):1993–5, apr 1992. ISSN 0008-5472.

23. S. David Nathanson and Lisa Nelson. Interstitial fluid pressure in breast cancer, benign breast conditions, and breast parenchyma. Annals of Surgical Oncology, 1 (4):333–338, jul 1994. ISSN 1068-9265. doi: 10.1007/BF03187139.

24. L.J. Liu and M. Schlesinger. MRI contrast agent concentration and tumor interstitial fluid pressure. Journal of Theoretical Biology, 406:52–60, oct 2016. ISSN 00225193. doi: 10.1016/j.jtbi.2016.06.027.

25. Long Jian Liu, Stephen L. Brown, James R. Ewing, Brigitte D. Ala, Kenneth M. Schneider, Mordechay Schlesinger, Long Jian Liu, Stephen L. Brown, James R. Ewing, Brigitte D. Ala, Kenneth M. Schneider, and Mordechay Schlesinger. Estimation of Tumor Interstitial Fluid Pressure (TIFP) Noninvasively. PLoS ONE, 11(7):e0140892, jul 2016. ISSN 1932-6203. doi: 10.1371/journal.pone.0140892.

26. Simon Walker-Samuel, Thomas A Roberts, Rajiv Ramasawmy, Jake S Burrell, Sean Peter Johnson, Bernard M Siow, Simon Richardson, Miguel R Gonçalves, Douglas Pendse, Simon P Robinson, R. Barbara Pedley, and Mark F Lythgoe. Investigating low-velocity fluid flow in tumors with convection-MRI. Cancer Research, 78(7):1859–1872, apr 2018. ISSN 15387445. doi: 10.1158/0008-5472.CAN-17-1546.

27. Paolo A. Netti, Laurence T. Baxter, Yves Boucher, Richard Skalak, and Rakesh K. Jain. Time-dependent Behavior of Interstitial Fluid Pressure in Solid Tumors: Implications for Drug Delivery. Cancer Res., 55(22):5451–5458, 1995. ISSN 0008-5472.

28. M Welter and H Rieger. Physical determinants of vascular network remodeling during tumor growth. Eur. Phys. J. E, 33:149–163, 2010. doi: 10.1140/epje/i2010-10611-6.

29. Vasileios Vavourakis, Peter A. Wijeratne, Rebecca Shipley, Marilena Loizidou, Triantafyllos Stylianopoulos, and David J. Hawkes. A Validated Multiscale In-Silico Model for Mechano-sensitive Tumour Angiogenesis and Growth. PLoS Computational Biology, 13(1):e1005259, jan 2017. ISSN 15537358. doi: 10.1371/journal.pcbi.1005259.

30. Triantafyllos Stylianopoulos, Konstantinos Soteriou, Dai Fukumura, and Rakesh K. Jain. Cationic Nanoparticles Have Superior Transvascular Flux into Solid Tumors: Insights from a Mathematical Model. Annals of Biomedical Engineering, 41(1):68–77, jan 2013. ISSN 0090-6964. doi: 10.1007/s10439-012-0630-4.

31. M Soltani and P Chen. Numerical Modeling of Interstitial Fluid Flow Coupled with Blood Flow through a Remodeled Solid Tumor Microvascular Network. PLoS ONE, 8(6):e67025, 2013. ISSN 19326203. doi: 10.1371/journal.pone.0067025.

32. M Sefidgar, M Soltani, K Raahemifar, M Sadeghi, H Bazmara, M Bazargan, and M Mousavi Naeenian. Numerical modeling of drug delivery in a dynamic solid tumor microvasculature. Microvascular Research, 99:43–56, 2015. ISSN 10959319. doi: 10.1016/j.mvr.2015.02.007.

33. M. Mohammadi and P. Chen. Effect of microvascular distribution and its density on interstitial fluid pressure in solid tumors: A computational model. Microvascular Research, 101:26–32, sep 2015. ISSN 0026-2862. doi: 10.1016/J.MVR.2015.06.001.

34. Michael Welter and Heiko Rieger. Interstitial Fluid Flow and Drug Delivery in Vascularized Tumors: A Computational Model. PLoS ONE, 8(8), 2013. ISSN 19326203. doi: 10.1371/journal.pone.0070395.

35. Jianbing Zhao, Howard Salmon, and Malisa Sarntinoranont. Effect of heterogeneous vasculature on interstitial transport within a solid tumor. Microvascular Research, 73(3):224–236, 2007. ISSN 00262862. doi: 10.1016/j.mvr.2006.12.003.

36. Gregory L. Pishko, Garrett W. Astary, Thomas H. Mareci, and Malisa Sarntinoranont. Sensitivity Analysis of an Image-Based Solid Tumor Computational Model with Heterogeneous Vasculature and Porosity. Annals of Biomedical Engineering, 39(9):2360–2373, sep 2011. ISSN 0090-6964. doi: 10.1007/s10439-011-0349-7.

37. Johnathon R Walls, John G Sled, James Sharpe, and R Mark Henkelman. Resolution improvement in emission optical projection tomography. Physics in Medicine and Biology, 52(10):2775–2790, may 2007. ISSN 0031-9155. doi: 10.1088/0031-9155/52/10/010.

38. Angela D’Esposito, Paul W. Sweeney, Morium Ali, Magdy Saleh, Rajiv Ramasawmy, Thomas A. Roberts, Giulia Agliardi, Adrien Desjardins, Mark F. Lythgoe, R. Barbara Pedley, Rebecca Shipley, and Simon Walker-Samuel. Computational fluid dynamics with imaging of cleared tissue and of in vivo perfusion predicts drug uptake and treatment responses in tumours. Nature Biomedical Engineering, 2(10):773–787, oct 2018. ISSN 2157-846X. doi: 10.1038/s41551-018-0306-y.

39. Brendan C. Fry, Jack Lee, Nicolas P. Smith, and Timothy W. Secomb. Estimation of Blood Flow Rates in Large Microvascular Networks. Microcirculation, 19: 530–538, 2012. ISSN 10739688. doi: 10.1111/j.1549-8719.2012.00184.x.

40. Timothy W. Secomb, Richard Hsu, Eric Y. H. Park, and Mark W. Dewhirst. Green’s Function Methods for Analysis of Oxygen Delivery to Tissue by Microvascular Networks. Annals of biomedical engineering, 32(11):1519–1529, 2004. ISSN 00906964. doi: 10.1114/B:ABME.0000049036.08817.44.

41. P Workman, E O Aboagye, F Balkwill, A Balmain, G Bruder, D J Chaplin, J A Double, J Everitt, D A H Farningham, M J Glennie, L R Kelland, V Robinson, I J Stratford, G M Tozer, S Watson, S R Wedge, S A Eccles, V Navaratnam, and S Ryder. Guidelines for the welfare and use of animals in cancer research. British Journal of Cancer, 102(11):1555–1577, may 2010. ISSN 0007-0920. doi: 10.1038/sj.bjc.6605642.

42. A. R. Pries, T. W. Secomb, P. Gaehtgens, and J. F. Gross. Blood flow in microvascular networks. Experiments and simulation. Circulation Research, 67(4): 826–834, oct 1990. ISSN 0009-7330. doi: 10.1161/01.RES.67.4.826.

43. A R Pries, T W Secomb, and P Gaehtgens. Structure and hemodynamics of microvascular networks: heterogeneity and correlations. The American journal of physiology, 269(5 Pt 2):H1713–22, nov 1995. ISSN 0002-9513.

44. Spyros k. Stamatelos, Eugene Kim, Arvind P. Pathak, Popel, and Aleksander S. A bioimage informatics based reconstruction of breast tumor microvasculature with computational blood flow predictions. Microvasc Res., 29(6):997–1003, 2012. ISSN 15378276. doi: 10.1016/j.biotechadv.2011.08.021.Secreted.

45. Paul W. Sweeney, Simon Walker-Samuel, and Rebecca J. Shipley. Insights into cerebral haemodynamics and oxygenation utilising in vivo mural cell imaging and mathematical modelling. Scientific Reports, 8(1):1373, dec 2018a. ISSN 20452322. doi: 10.1038/s41598-017-19086-z.

46. Zahra Farid, Amani H. Saleem, Baraa K. Al-Khazraji, Dwayne N. Jackson, and Daniel Goldman. Estimating blood flow in skeletal muscle arteriolar trees reconstructed from in vivo data using the Fry approach. Microcirculation, 24(5), 2017. ISSN 15498719. doi: 10.1111/micc.12378.

47. Timothy W Secomb. A Green’s function method for simulation of time-dependent solute transport and reaction in realistic microvascular geometries. Mathematical Medicine and Biology, 33(4):475–494, oct 2016. ISSN 14778602. doi: 10.1093/imammb/dqv031.

48. A R Pries and T W Secomb. Microvascular blood viscosity in vivo and the endothelial surface layer. American journal of physiology. Heart and circulatory physiology, 289(6):H2657–H2664, 2005. ISSN 0363-6135. doi: 10.1152/ajpheart.00297.2005.

49. A R Pries, K Ley, M Claassen, and P Gaehtgens. Red cell distribution at microvascular bifurcations. Microvascular research, 38(1):81–101, jul 1989. ISSN 0026-2862.

50. A R Pries, K Ley, and P Gaehtgens. Generalization of the Fahraeus principle for microvessel networks. The American journal of physiology, 251(6 Pt 2):H1324–32, dec 1986. ISSN 0002-9513.

51. Axel R. Pries and Timothy W. Secomb. Blood Flow in Microvascular Networks. In Comprehensive Physiology, pages 3–36. John Wiley & Sons, Inc., Hoboken, NJ, USA, jan 2011. ISBN 9780470650714. doi: 10.1002/cphy.cp020401.

52. Laurence T Baxter and Rakesh K Jain. Transport of fluid and macromolecules in tumors. II. Role of heterogeneous perfusion and lymphatics. Microvascular Research, 40(2):246–263, sep 1990. ISSN 10959319. doi: 10.1016/0026-2862(90)90023-K.

53. Rakesh K. Jain. Normalization of Tumor Vasculature: An Emerging Concept in Antiangiogenic Therapy. Science, 307(5706), 2005.

54. Triantafyllos Stylianopoulos and Rakesh K Jain. Combining two strategies to improve perfusion and drug delivery in solid tumors. Proceedings of the National Academy of Sciences, 110(46):18632–18637, nov 2013. ISSN 0027-8424. doi: 10.1073/pnas.1318415110.

55. Paul William Sweeney, Simon Walker-Samuel, and Rebecca J Shipley. Vascular and interstitial flow solver for discrete microvascular networks, 2018b.

56. Timothy W. Secomb. Green’s function method for simulation of oxygen transport to tissue. http://physiology.arizona.edu/people/secomb/greens, 2011.

57. Conrad Sanderson and Ryan Curtin. Armadillo: a template-based C++ library for linear algebra. Journal of Open Source Software, Vol. 1:p.26, 2016.

58. Jon-Vidar Gaustad, Trude G Simonsen, Kjetil G Brurberg, Else Marie Huuse, and Einar K Rofstad. Blood supply in melanoma xenografts is governed by the morphology of the supplying arteries. Neoplasia (New York, N.Y.), 11(3):277–85, mar 2009. ISSN 1476-5586.

59. Shunichi Morikawa, Peter Baluk, Toshiyuki Kaidoh, Amy Haskell, Rakesh K Jain, and Donald M McDonald. Abnormalities in pericytes on blood vessels and endothelial sprouts in tumors. The American journal of pathology, 160(3): 985–1000, mar 2002. ISSN 0002-9440. doi: 10.1016/S0002-9440(10)64920-6.

60. FE Curry. Mechanics and thermodynamics of transcapillary exchange, Handbook of Physiology, Section 2: The Cardiovascular System. jan 1984.

61. R. A. Brace and A. C. Guyton. Transmission of Applied Pressure Through Tissues: Interstitial Fluid Pressure, Solid Tissue Pressure, and Total Tissue Pressure. Experimental Biology and Medicine, 154(2):164–167, feb 1977. ISSN 1535-3702. doi: 10.3181/00379727-154-39628.

62. Chang-Geol Lee, Marcus Heijn, Emmanuelle di Tomaso, Genevieve Griffon-Etienne, M. Ancukiewicz, Chieko Koike, K. R. Park, Napoleone Ferrara, Rakesh K. Jain, Herman D. Suit, and Yves Boucher. Anti-Vascular Endothelial Growth Factor Treatment Augments Tumor Radiation Response under Normoxic or Hypoxic Conditions. Cancer Research, 60(19), 2000.

63. M. Soltani and P. Chen. Numerical modeling of fluid flow in solid tumors. PLoS ONE, 6(6):e20344, jun 2011. ISSN 19326203. doi: 10.1371/journal.pone.0020344.

64. Eugene Kim, Spyros Stamatelos, Jana Cebulla, Zaver M. Bhujwalla, Aleksander S. Popel, and Arvind P. Pathak. Multiscale imaging and computational modeling of blood flow in the tumor vasculature. Annals of Biomedical Engineering, 40(11): 2425–2441, 2012. ISSN 00906964. doi: 10.1007/s10439-012-0585-5.

65. Ethaar El-Emir, Geoffrey M. Boxer, Ingrid A. Petrie, Robert W. Boden, Jason L.J. Dearling, Richard H.J. Begent, and R. Barbara Pedley. Tumour parameters affected by combretastatin A-4 phosphate therapy in a human colorectal xenograft model in nude mice. European Journal of Cancer, 41(5): 799–806, mar 2005. ISSN 09598049. doi: 10.1016/j.ejca.2005.01.001.

66. Trude G. Simonsen, Jon-Vidar Gaustad, Marit N. Leinaas, and Einar K. Rofstad. High Interstitial Fluid Pressure Is Associated with Tumor-Line Specific Vascular Abnormalities in Human Melanoma Xenografts. PLoS ONE, 7(6):e40006, jun 2012. ISSN 1932-6203. doi: 10.1371/journal.pone.0040006.

67. Rakesh K. Jain, Ricky T. Tong, and Lance L. Munn. Effect of vascular normalization by antiangiogenic therapy on interstitial hypertension, peritumor edema, and lymphatic metastasis: Insights from a mathematical model. Cancer Research, 67(6):2729–2735, mar 2007. ISSN 00085472. doi: 10.1158/0008-5472.CAN-06-4102.

68. Vikash P. Chauhan, Triantafyllos Stylianopoulos, John D. Martin, Zoran Popovi Ä, Ou Chen, Walid S. Kamoun, Moungi G. Bawendi, Dai Fukumura, and Rakesh K. Jain. Normalization of tumour blood vessels improves the delivery of nanomedicines in a size-dependent manner. Nature Nanotechnology, 7(6):383–388, jun 2012. ISSN 17483395. doi: 10.1038/nnano.2012.45.

69. Hadi T. Nia, Hao Liu, Giorgio Seano, Meenal Datta, Dennis Jones, Nuh Rahbari, Joao Incio, Vikash P. Chauhan, Keehoon Jung, John D. Martin, Vasileios Askoxylakis, Timothy P. Padera, Dai Fukumura, Yves Boucher, Francis J. Hornicek, Alan J. Grodzinsky, James W. Baish, Lance L. Munn, and Rakesh K. Jain. Solid stress and elastic energy as measures of tumour mechanopathology. Nature Biomedical Engineering, 1(1), 2017. ISSN 2157846X. doi: 10.1038/s41551-016-0004.

